# Vascular draining confounds laminar decoding in fMRI

**DOI:** 10.1101/2025.08.26.672278

**Authors:** Jonas Karolis Degutis, Denis Chaimow, Romy Lorenz

## Abstract

Laminar fMRI using GE-BOLD is vulnerable to spatial blurring from intracortical veins, while multivariate pattern analysis (MVPA) is often assumed to mitigate these biases. Yet, this assumption has not been systematically investigated. We thus developed a mechanistic laminar response model that simulates voxel-wise patterns across cortical depths, incorporating a vascular draining model. We conducted simulations in which the ground-truth signal originated in a single, several, or across all layers, and applied standard MVPA decoding before and after deconvolution of the draining effect. Decoding accuracies were consistently influenced by draining veins: deep-origin signals yielded above-chance decoding in superficial layers, and null scenarios produced false positives in middle or deep layers. Vascular deconvolution enhanced specificity in single-layer cases but did not resolve ambiguities in null decoding profiles. Simulating six thinner layers improved decoding accuracies, especially in the deconvolved signal scenarios. These findings demonstrate that multivariate techniques are not inherently immune to vascular biases, but also demonstrate that careful modeling can help correct draining effects.

## Introduction

Cortical gray matter layers are a fundamental organizational principle of the neocortex, characterized by their distinct connections and functions. Ultra-high-field fMRI enables investigations of layer-specific functions in humans by measuring differences between laminar activations in both health (Lawrence, Formisano, et al., 2019; Yang et al., 2021) and disease (Haarsma et al., 2022). As such, laminar fMRI has contributed to understanding the interplay between feedforward and feedback signals in sensory areas (Aitken et al., 2020; Kok et al., 2016; Lawrence et al., 2018), as well as the localization of cognitive functions within association cortices at the mesoscopic level (Chaimow et al., 2025; Degutis et al., 2024; Sharoh et al., 2019). Many laminar fMRI studies use univariate methods to compare activation levels between cortical layers to draw conclusions about layer-specific processing. These approaches have been applied to a range of cortical regions, including early visual cortices (Aitken et al., 2020; de Hollander et al., 2021; Iamshchinina, Kaiser, et al., 2021; Kok et al., 2016; Koopmans et al., 2010; Lawrence et al., 2018; Lawrence, Norris, et al., 2019; Thomas et al., 2024), auditory regions (De Martino et al., 2015), primary motor area (Huber et al., 2017), and frontal cortices (Chaimow et al., 2025; Degutis et al., 2024).

Even with the development of alternative fMRI sequences (Huber et al., 2019; Iamshchinina, Haenelt, et al., 2021; Norris, 2012; Ramadan et al., 2025), most studies to date rely on GE-BOLD imaging, because of its higher sensitivity. However, it is well established that GE-BOLD suffers from reduced spatial specificity, as the signal is dominated by draining intracortical veins that run perpendicularly to the surface and drain blood from deeper layers towards the pial surface (Duvernoy et al., 1981). The signal from more superficial layers is thus blurred by signals originating in deeper layers (Huber et al., 2019; Koopmans et al., 2010). As a result of this leakage, superficial layers show a stronger, but mixed signal, compromising the spatial specificity of laminar findings in univariate analyses.

Some studies aim to mitigate the draining vein effect by subtracting experimental conditions, which removes non-specific layer effects shared by both conditions (Aitken et al., 2020; Degutis et al., 2024; Kok et al., 2016; Lawrence et al., 2018; Lawrence, Norris, et al., 2019) or use neurophysiologically inspired depth-dependent BOLD response models based on hemodynamics and vasculature (Havlicek & Uludağ, 2020; Heinzle et al., 2016; Markuerkiaga et al., 2016, 2021; Marquardt et al., 2018; Uludag & Havlicek, 2021) to deconvolve the mixed signal to isolate layer-specific univariate responses.

Interestingly, multivariate approaches, which capture distributed patterns of voxel activity within individual layers reflecting distinct information processing, are also often assumed to mitigate these biases. The rationale being that decoding performance remains stable despite univariate superficial bias, provided that both the effect and the noise scale proportionally with overall signal strength (Huang et al., 2021). Studies employing these approaches have reported higher decoding accuracies in deep and middle layers compared to superficial layers, suggesting that these methods better capture layer-specific neural processes and reveal the fine-scaled nature of decoded representations (Vizioli et al., 2020) without the prior need to address draining effects. Multivariate encoding models have shown higher deep and middle layer decoding accuracies revealing layer-specific thalamocortical input to V1 (de Hollander et al., 2021). Multivariate pattern analyses have also been used to distinguish between deep and superficial layer feedback of imagery and illusory content (Bergmann et al., 2024), expected and unexpected orientation stimuli in V1 (Thomas et al., 2024), and storage of rotated working memory stimuli in deep and superficial feedback layers of V1 (Iamshchinina, Kaiser, et al., 2021).

Despite the growing application of multivariate techniques to laminar fMRI (Bergmann et al., 2024; Degutis et al., 2024; Thomas et al., 2024), the extent to which these methods are affected by vascular signal leakage has not been systematically investigated. Understanding this is crucial, as multivariate techniques not only detect the presence of neural activation but can also reveal the representational content of signals. This opens a unique opportunity to directly relate human fMRI measurements to the rich body of laminar circuit models derived from invasive animal studies (Bastos et al., 2012, 2018; Felleman & Van Essen, 1991; Mendoza-Halliday et al., 2024; van Kerkoerle et al., 2017), where the anatomical origin and functional role of layer-specific signals can be determined with high precision. Demonstrating that fMRI-based multivariate analyses can reliably localize and interpret layer-specific representations would strengthen the translational bridge between human mesoscopic imaging and mechanistic insights from animal neurophysiology. Conversely, if vascular biases distort these patterns, interpretations about the laminar origin of specific information could be fundamentally compromised.

Here, we address this gap by developing a mechanistic laminar response model that simulates multivariate patterns across cortical depth while incorporating vascular biases. Using this model, we performed exhaustive simulations to assess the impact of vascular draining effects on multivariate analyses of layer-specific GE-BOLD responses. More precisely, we ran multivariate pattern analyses on simulated responses where the ground-truth signal originated from: (i) a single layer, (ii) two layers, or (iii) all three layers at once. If multivariate decoding were subject to the same bias as univariate methods, we would expect them to yield inaccurate layer responses in all scenarios. Additionally, given that cortical regions differ in baseline decoding accuracy (Bhandari et al., 2018) as a possible result of difference in their neurobiology, like the size of columnar structures (Goldman-Rakic, 1995), we also examined whether decoding accuracy depends on an interaction between the spatial scale of the patterns and cortical layer. Finally, we also simulated six rather than three layers to see whether this can provide more accurate laminar decoding accuracy profiles.

## Methods

### Depth-dependent multivariate pattern simulation model

We built a mechanistic model to simulate cortical multivariate patterns across different depths of the cortex following previous work (Chaimow, Uğurbil, et al., 2018). As in the previous model, we included four main stages in the simulation: the generation of the multivariate pattern and its neuronal response, spatial BOLD activation, and MR voxel sampling. We extended the simulations from two-dimensions to the third depth (or laminar) dimension.

Per simulation, we generated two types of conditions represented by multivariate activation patterns. Each simulation had 50 trials per condition, which constituted a single simulated dataset. We simulated a total of 20 datasets per simulation type.

For each simulation, the initial multivariate pattern was generated by the spatial filtering of Gaussian white noise (Rojer & Schwartz, 1990) and was parametrized by the main frequency *ρ*; the inverse of *ρ* indicated the pattern’s cycle length or twice the length of a column. Additionally, an irregularity parameter δ/ρ controls the width of the distribution of underlying spatial frequency components. As part of the simulations, we systematically varied *ρ* to study the effect of spatial frequency (i.e., the granularity of the multivariate response pattern) on decoding accuracies. The multivariate pattern simulation grid had the dimensions *N* x *N* x *N_Depth_*. For each depth we simulated a columnar grid generated based on a predefined seed value.

Each of the multivariate neural patterns were differential neuronal responses, modeled as the difference between two orthogonal neuronal responses that can be seen as a difference between two conditions. The differential neuronal response from the two responses was normalized resulting in a spatial average of the pattern that was equal to the BOLD amplitude parameter *β*, which was set to 3.5%, an expected response in GE-BOLD imaging at 7T (Chaimow, Uğurbil, et al., 2018). Additionally, a baseline neuronal activation *b* was added to not have any negative values due to the fluctuation of responses.

The BOLD response was simulated by convolving the neuronal activity patterns with a point-spread function of a predefined FWHM (Chaimow, Yacoub, et al., 2018; Shmuel et al., 2007). The FWHM increased from deeper to superficial depths with values ranging from 0.83 to 1.78 mm (Fracasso et al., 2021). We additionally smoothed the signal in the depth dimension by applying a point-spread function of the same FWHM as in-plane. A vascular model was then applied to induce the draining of the signal from deep to superficial layers. We applied the model proposed by (Markuerkiaga et al., 2016, 2021); multiplying the laminar responses with a lower triangle matrix consisting of peak-to-tail values blurred all signals from the second-to-deepest depth by adding a proportion of the signal originating from all previous depths. This provided a univariate increase in the simulated BOLD signal and blurred the multivariate signal.

The BOLD signal across the three dimensions was MR-sampled using a three-dimensional Fourier transform to downsample and then reconstruct the signal with an isotropic voxel size of 1 mm^3^ thus also inducing partial voluming effects. To avoid a wrap-around effect between the superficial and deep depths, the BOLD signal matrix was zero-padded in the depth dimension.

Thermal and physiological noise was added to the reconstructed MR sampled voxel responses by using a 7T-specific noise model (Chaimow, Uğurbil, et al., 2018; Triantafyllou et al., 2005). Assuming the multivariate patterns are based on a single volume per trial, we did not include temporal correlations of physiological noise. However, physiological noise amplitudes were scaled for each depth with higher noise levels in the middle layers (Koopmans et al., 2011).

### Types of simulations

We ran seven main types of simulations: three simulations where the origin of the signal was either the superficial, middle, or deep layers; two simulations where the signal was in superficial and deep layers (later referred to as top-down); and two null simulations where all layers had signals.

In our primary simulation, we created two multivariate patterns that were present across all simulated trials in specific simulated depths. We ran three of these simulations with the multivariate signal originating either from the deep, middle, or superficial depths. All other depths had multivariate patterns with the same parameters except they did not repeat across trials (Fig. 1b). We also simulated two null scenarios. In the first, all depths had the same two multivariate patterns repeating across trials (Fig. 3a), while in the second, each of the three layers had two repeating patterns, but the pairs differed in each layer (Fig. 3e). We simulated two scenarios where deep and superficial layers either had the same or different pairs of patterns (Fig. 2).

We also ran a voxel misalignment analysis to reproduce real-life scenarios of mis-segmentation of gray matter layers, CSF, and white matter. The analysis was performed on the simulated BOLD data of the aforementioned simulations. We randomly selected a certain percentage of voxels in each layer. These voxels were of the same indices in each layer, thus simulating a scenario where voxels from the same column are misaligned. The selected voxels were then switched between adjacent layers (e.g. deep-middle, middle-superficial). Superficial voxels were also switched with simulated CSF, while deep voxels were also switched with simulated white matter. CSF and white matter were simulated as white-noise with different scaling factors with their values being similar to either the most superficial or deep depths, respectively (Koopmans et al., 2011). This analysis was run for each of the 20 iterations of the simulations and additionally for percentages ranging from 1% -40% misaligned voxels. We then computed t-values of the decoding accuracy between a layer-of-interest (where the multivariate signal originated) and the other two layers to see whether decoding remained stable across increasing misalignment.

To see whether decoding differences between layers can be enhanced, we applied deconvolution on the simulated laminar BOLD signal, a technique usually only used for univariate analyses. We inverted the draining vein vascular model matrix and used this to deconvolve the laminar BOLD signal. We applied deconvolution to all five of the simulation types. Since draining is dependent on local vasculature, in a further analysis the parameters of the draining vein model were changed to either increase or decrease the draining vein strength. We then used the initial parameters to invert the matrix. This also tested a more realistic scenario where the exact parameters of draining strength are unknown.

### Data availability statement

The code for the model and corresponding data are archived on Zenodo: 10.5281/zenodo.16947106.

## Results

**Figure 1.**
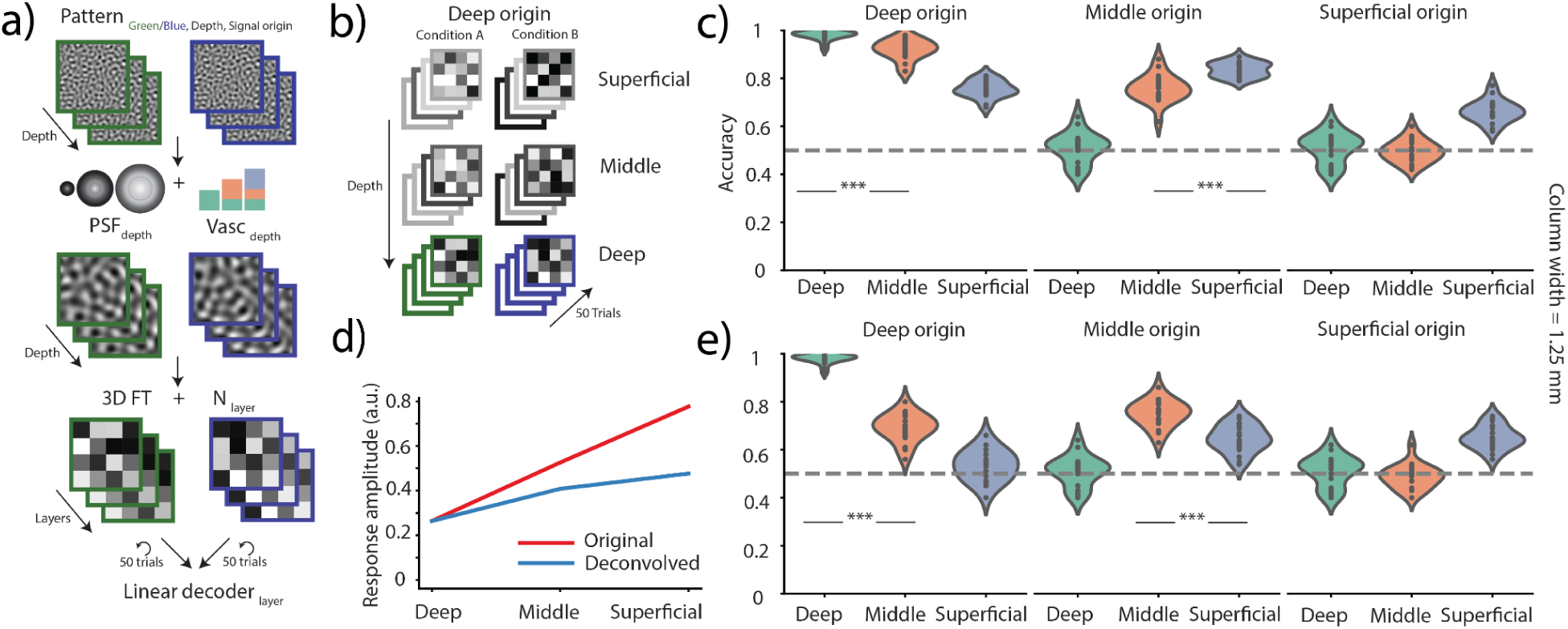
Model and single layer origin simulations. **a)** The simulation model. Multivariate patterns were created of a given spatial frequency and convolved with a BOLD depth-dependent point-spread function for each laminar depth. A vascular model was applied across depth draining the signal from deep to superficial depths. Voxel responses were MR sampled into three layers using a 3D Fourier transform (FT) and depth-dependent thermal noise added. A total of 50 trials were generated of each pattern per iteration of a simulation. A linear decoder (SVM) was then used to classify the two patterns (green and blue) independently in each layer. **b)** Deep origin simulation. Multivariate patterns were only consistent in deep layers (green and blue), while other layers had different multivariate patterns in each of the 50 trials of each condition. **c)** Decoding accuracies across layers. Multivariate patterns were simulated as indicated in b), but they were consistent across trials only in certain layers (e.g. deep, middle or superficial origin). Pairwise comparisons are only done between the layer of origin and the adjacent superficial layer. **d)** Univariate response across layers of the original and deconvolved signals. Each point within the violin plot indicates one iteration (total of 20) of the simulation. **e)** Same as c) but the voxel responses were deconvolved using an inversion of the vascular model with the same parameters as the initial convolution. *** *p* < 0.001.

### Draining of multivariate patterns results in decoding from layers of signal origin and non-origin

We simulated cortical multivariate patterns across different depths of the cortex. Our simulation involved four stages: generating multivariate patterns, modeling neuronal responses, simulating spatial BOLD activation, and MR voxel sampling (Fig. 1a). The quasi-periodic spatial multivariate patterns were created with a given spatial frequency in each depth and were then convolved with depth-varying BOLD point-spread functions followed by a vascular model that drained the signal from deep toward superficial layers (Markuerkiaga et al., 2016, 2021). The BOLD signal was then MR sampled, and layer-dependent noise added. In all analyses, two distinguishable multivariate patterns were generated with the origin of these patterns varying depending on the analysis. A linear classifier was then used to decode these patterns from each layer irrespective of the origin of the multivariate signal. Most laminar decoding studies infer layer specificity by comparing decoding accuracies across layers. We adopt the same approach: in each simulation, we compare decoding accuracy differences between layers, and since we know the origin of the signal we can explicitly flag inaccurate results.

The primary analysis simulated pairs of distinguishable patterns originating from one of the three layers: deep, middle, or superficial (Fig. 1b; depicts the deep origin simulation). In the case that the signal originated in deep depths, the decoding accuracies were above-chance in all layers and showed significant variation (one-way ANOVA, *F*(2, 17) = 236, *p* < 0.001; Fig. 1c, left subpanel), driven by a robust difference between the deep and middle layers (paired-sample t-test, *t*(19) = 5.95, *p* < 0.001). The deep layer decoding accuracy was close to ceiling, likely because the underlying multivariate signal incurred less blurring (see Methods on point-spread function), preserving the spatial size of the patterns, and had lower depth-dependent added noise (Koopmans et al., 2011). However, in this simulation we see above-chance decoding in both the middle and superficial layers; this indicates that, despite the assumption of multivariate analyses being more spatially specific, we still observe some draining of the multivariate pattern from deep to superficial layers.

**Figure 2.**
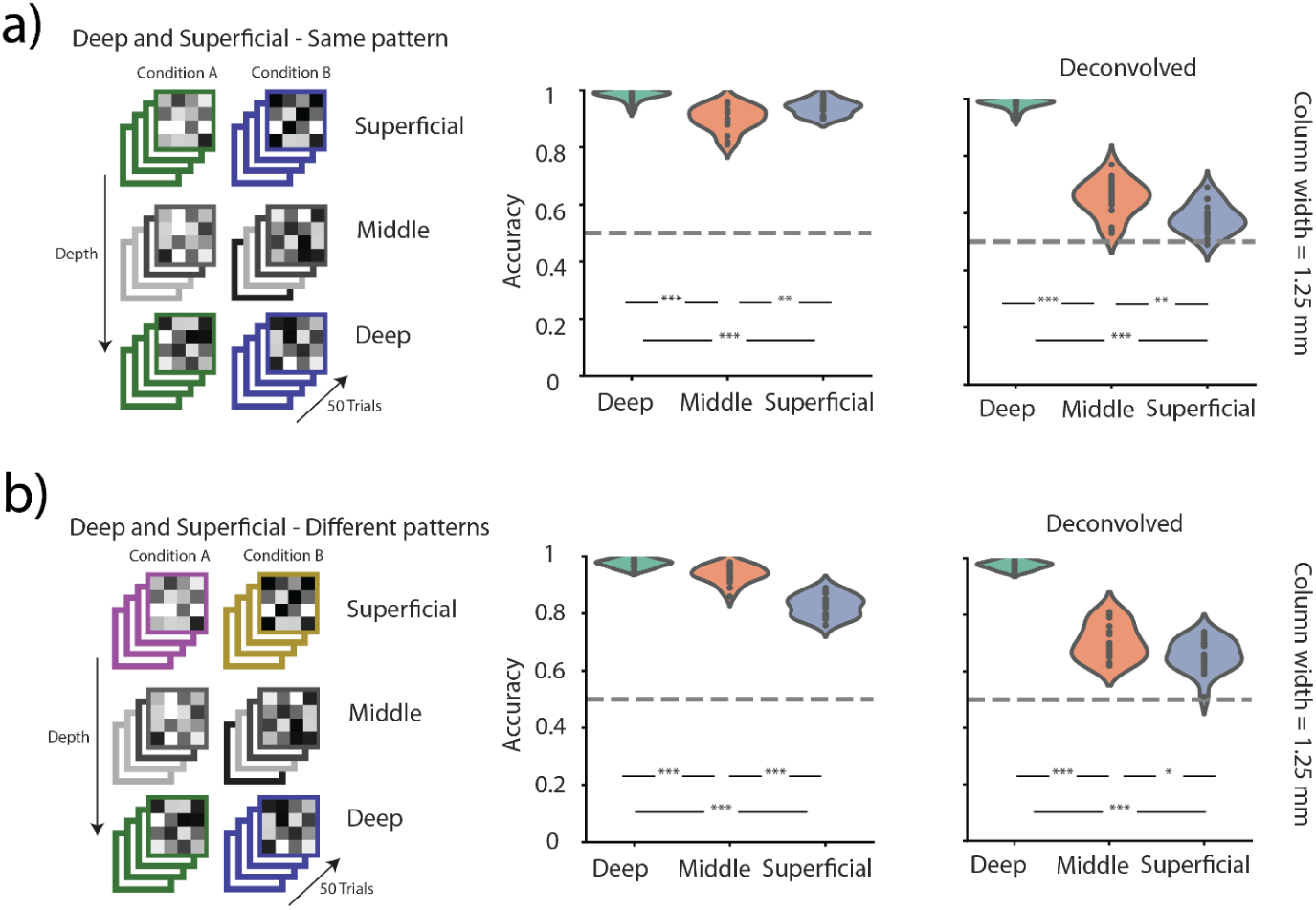
Top-down scenarios. **a)** Deep and superficial origin simulation containing the same patterns in both layers. Each point within the violin plot indicates one iteration (total of 20) of the simulation. Left and right show decoding accuracies on non-deconvolved and deconvolved layer signals, respectively. **b)** Same as a), but the pattern pairs in the superficial and deep layers are different. * *p* < 0.05, ** *p* < 0.01, *** *p* < 0.001.

Laminar decoding accuracies were significantly different between layers when the signal originated from the middle depths (one-way ANOVA, *F*(2, 17) = 219, *p* < 0.001; Fig. 1c, middle subpanel), but this was driven by higher superficial rather than middle decoding accuracy (paired-sample t-test, *t*(19) = 5.84, *p* < 0.001). In such cases, one might erroneously conclude that superficial layers contribute more despite the signal originating from middle layers.

Finally, when the underlying signal originated from superficial depths, only the superficial layers had above-chance accuracy (one-way ANOVA, *F*(2, 17) = 63.7, *p* < 0.001); however, due to the broader point-spread function resulting in a blurred pattern, the decoding accuracy was lower compared to simulations of deep and middle signal origins (Fig. 1c, right subpanel). Overall, we find that the layer of origin showed maximal decoding in the deep and superficial origin relative to other layers; however, vascular draining resulted in considerable spread of decodable information.

One way of overcoming the draining vein effect in GE-BOLD imaging is by deconvolving the laminar signal (de Hollander et al., 2021; Markuerkiaga et al., 2021), yet this has primarily been applied to univariate analyses, and the effects of deconvolution on multivariate patterns have not been examined. We aimed to remove the draining bias induced by our vascular model deconvolving the signal in each layer (see Methods). We find increased precision in the decoding accuracies in all single-origin scenarios: a large difference between deep and middle layers when the signal originated from deep depths (paired-sample t-test, *t*(19) = 19.5, *p* < 0.001; Fig. 1e, left), a reversal in decoding accuracy difference with higher middle layer decoding compared to superficial layers when the signal originated from middle depths (paired-sample t-test, *t*(19) = 4.62, *p* < 0.001; Fig. 1e, middle), and the same accuracy profile when the signal originated from superficial layers (Fig. 1e, right). The deconvolution model also reduced the univariate amplitude of the middle and superficial layers, but a bias towards superficial layers remained (Fig. 1d).

### Simulations of a canonical top-down signal result in mixed precision

Next, we simulated a canonical top-down feedback scenario where signals originate in the deep and superficial layers (Iamshchinina, Kaiser, et al., 2021; Lawrence et al., 2018; Lawrence, Norris, et al., 2019; Thomas et al., 2024). In one scenario the patterns in the deep and superficial layers were the same - reflecting a shared feedback population response - (Fig. 2a, left panel) and resulted in highest decoding accuracies in the deep and superficial layers (Figure 2a, middle panel; paired-sample t-test, deep-middle: *t*(19) = 7.78, *p* < 0.001; middle-superficial: *t*(19) = 3.78, *p* = 0.0013). Unlike in the single-layer origin scenarios, deconvolution failed to improve the precision of the decoding accuracy and rather induced false positives, resulting in higher middle rather than superficial layer accuracy (*t*(19) = 3.13, *p* = 0.006). We also simulated a scenario where the signal originated from the deep and superficial layers, but the pairs of patterns were different in each layer - indicating distinct layer-specific population responses (Fig. 2b, left panel). In both the non-deconvolved and deconvolved scenarios we found a decreasing decoding accuracy trend from deep towards superficial layers, which might lead to a false conclusion of a higher population response in deep layers.

**Figure 3.**
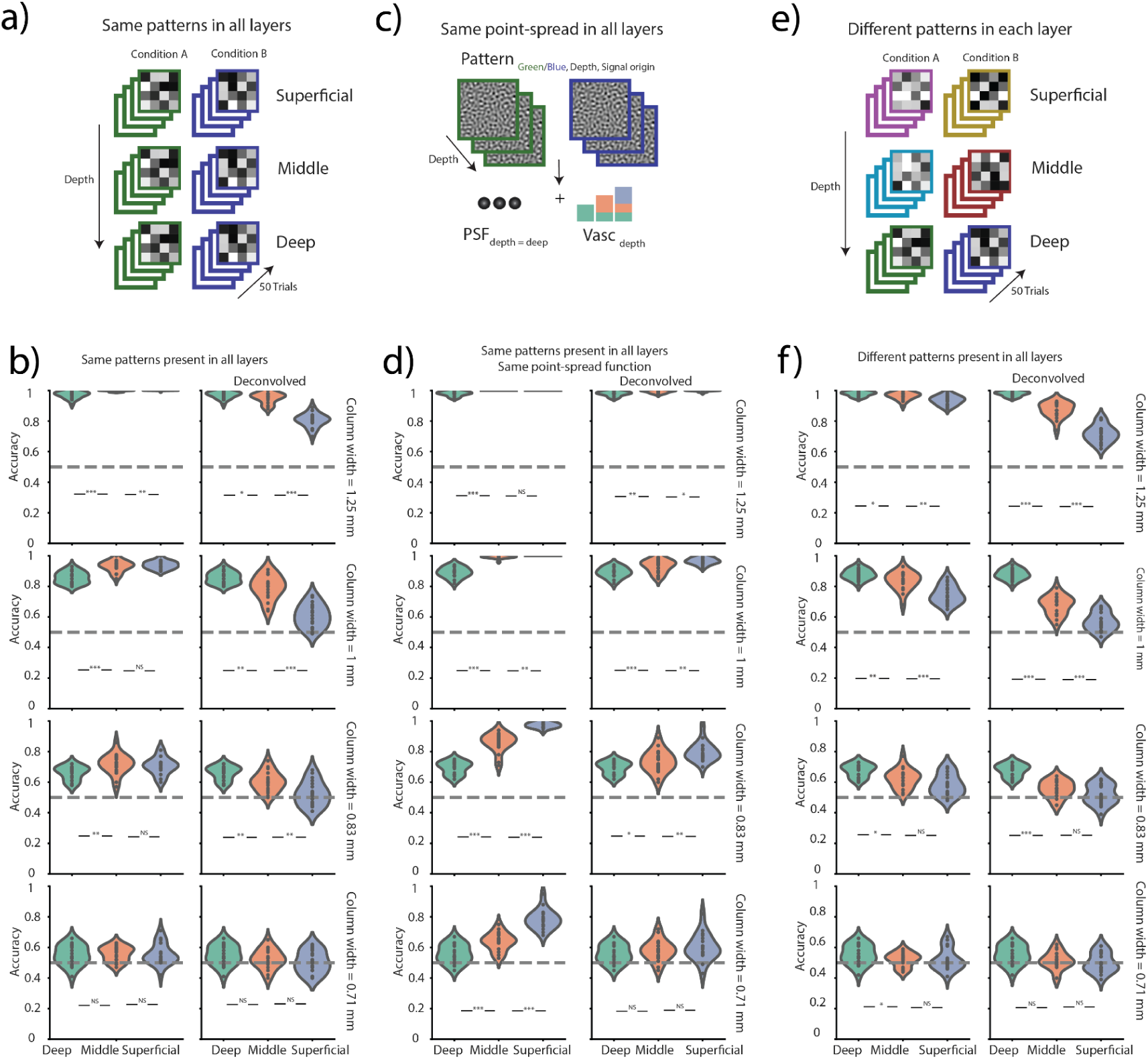
Null scenarios. **a)** Illustration of the same pattern simulation. Two distinguishable patterns in all layers. **b)** Decoding across different spatial scales of multivariate patterns. Each point within the violin plot indicates one iteration (total of 20) of the simulation. Left and right show decoding accuracies on non-deconvolved and deconvolved layer signals, respectively. **c)** Illustration of the process of obtaining the same point-spread function in each layer. **d)** Same as b), but the point-spread function does not vary across depth (FWHM = 0.83 mm). **e)** Illustration of the different pattern simulation. Two distinguishable patterns in each layer. **f)** Same as b), but for different patterns. * *p* < 0.05, ** *p* < 0.01, *** *p* < 0.001.

### Same pattern null scenario results in false positives in middle layer

We further explored non-layer-specific null scenarios where we simulated either the activation of all layers by the same two multivariate patterns (Fig. 3a) or by different pairs of patterns in all layers (Fig. 3e). The former simulation resembles layer-agnostic columnar processing, while the latter simulates different layer-specific responses to the same condition. In these simulations we also varied the spatial scale of patterns to see whether they might interact with decoding accuracies in the three layers. One might expect that a larger point-spread function in the superficial layer would blur smaller patterns resulting in lower decoding accuracies.

In the first scenario where all layers had the same two patterns present, we find that the spatial scale of the columnar pattern impacts layer-specificity of decoding results with lower decoding across all layers with decreasing spatial size of the multivariate columns (two-way ANOVA, main effect of spatial scale *F*(3, 228) = 1153, *p* < 0.001; Fig. 3b left). Additionally, we see layer-specific differences in decoding accuracy despite patterns being present in all layers (two-way ANOVA, main effect of layer *F*(2, 228) = 21.6, *p* < 0.001). There was also an interaction effect between columnar size and layers (*F*(6, 228) = 3.34, *p* = 0.0034; Fig. 3b, left), possibly driven by the decreasing difference between superficial and middle layer decoding accuracy: we see a significant difference only in the first largest columnar size scenario (see Supplementary Table 1). Overall, this indicates that in a null-scenario we get layer-specific decoding accuracies, i.e. false positives.

As in the previous simulation scenarios, we applied deconvolution with the aim of removing any laminar differences in decoding accuracy as per expectation in a null scenario. Similarly to the same-pattern top-down origin scenario (Fig. 2a), deconvolution reversed the decoding accuracy difference resulting in highest accuracies in the deep and lowest in the superficial layers (two-way ANOVA main effect of layer *F*(2, 228) = 134, *p* < 0.001; Fig. 3b, right), as a result of signal removal from the latter two layers; again leading to false positives. While this null scenario resulted in higher decoding accuracies in the superficial layers, the difference between superficial and middle was only significant for the largest columnar size. To understand why there might be an interaction between columnar size and layers, we tested whether a constant point-spread function across all layers would solidify the difference between middle and superficial depths in all spatial pattern sizes, as the spatial blurring of the multivariate patterns in each layer would stay constant. We thus ran the same null scenario analysis and kept the point-spread function fixed to the lowest value of 0.83 mm across all layers (Fig. 3c). As predicted, we find the highest decoding accuracies in superficial layers which remain significantly different compared to middle layers in most spatial sizes (see Supplementary Table 2 for paired t-tests). Deconvolution applied to this simulation resulted in a similar pattern of results. Intriguingly, these scenarios show that a larger point-spread function can impact the decoding of smaller-sized patterns.

### Different pattern null scenario yields layer-specific decoding in deep layer

In the second null scenario, we simulated different pairs of distinguishable patterns in each layer (Fig. 3e). Here, we might expect a blurring of patterns towards superficial layers both because the patterns originating in those layers have a larger point-spread, and because of the drainage of different patterns from deep and middle layers. As expected, we observe that decoding accuracy decreases from deep to superficial layers and accuracy across all layers decreases with smaller pattern sizes (Fig. 3f, left; see Supplementary Table 3 for paired t-tests). We deconvolved the layers resulting in a larger difference between deep, middle, and superficial layer decoding accuracies (Supplementary Table 3). We find higher decoding in deep layers (i.e. false positives) even though, again, the ground truth did not favor any layer.

**Figure 4:**
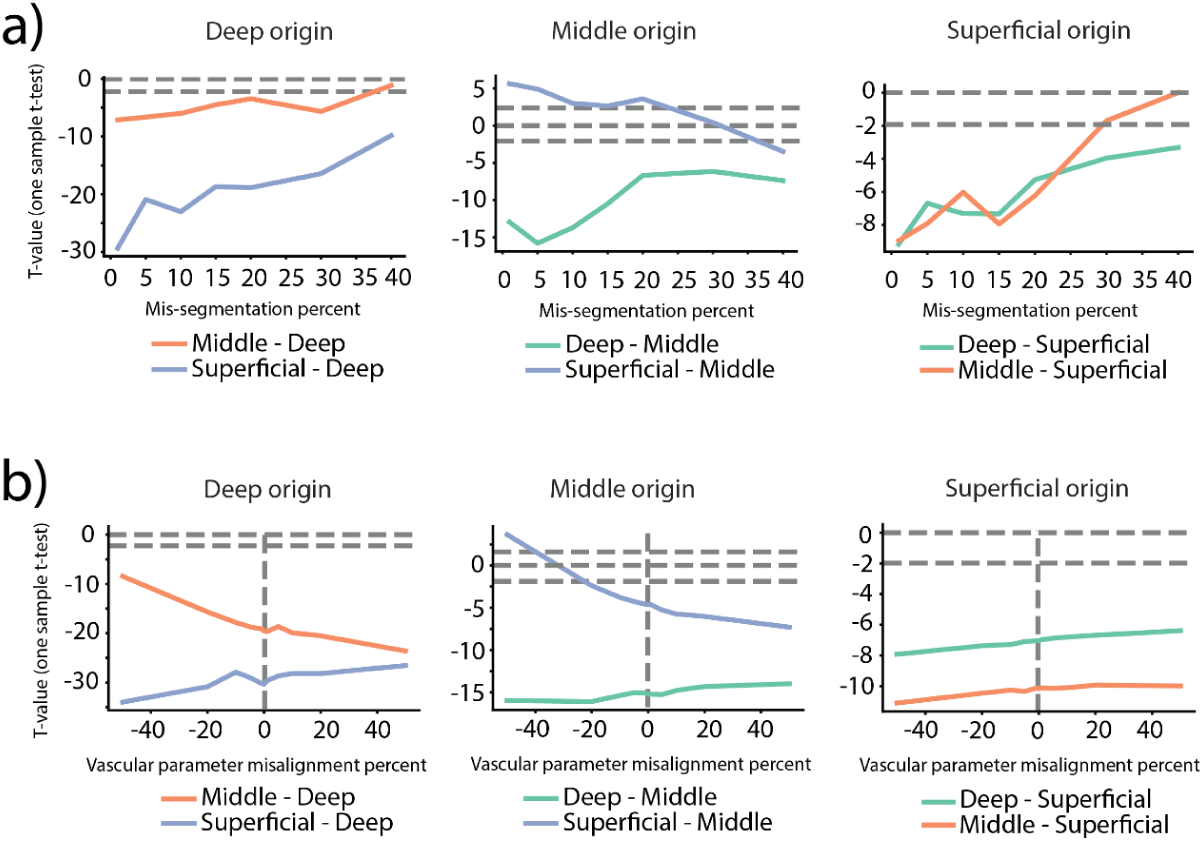
Stability of decoding. **a)** Mis-segmentation of voxels in the 1.25 mm columnar width simulations. Percentages of voxels from adjacent layers were switched in each trial and the decoding analysis was run on the new data. T-values compare decoding accuracy between each pair of layers. **b)** Same as c), but for vascular parameter misalignment. Misalignment indicates the percent difference between the draining peak-to-tail parameter used to convolve and deconvolve the signal. A negative percentage indicates a smaller value used for the convolution, while a positive value indicates a larger value.

### Layer mis-segmentation preserves decoding accuracy contrast

Laminar studies use (semi-)manual (Huber et al., 2017) or automatic (Chaimow et al., 2025; Degutis et al., 2024) preprocessing steps to segment the cortex and delineate the layers. Since the main simulations assumed unrealistic perfect laminar delineation, which one would not expect even in hand-crafted segmentations, we tested the effect of layer mis-segmentation on decoding results. We simulate the inaccuracies in layer delineation that can arise from segmentation methods. Specifically, we selected a percent of voxel responses (non-deconvolved signal) from each of the three layers and swapped them with voxels from an adjacent layer (e.g. deep voxels switched with middle voxels and middle voxels switched with superficial voxels) or CSF or white matter in the case of superficial and deep, respectively. We plot the t-values comparing decoding accuracies from each layer-of-interest to the other two layers in the three single-layer origin scenarios (Fig. 4a).

When the signal originates from deep depths, we find the sign of the t-values comparing superficial and middle layers to deep remains stable with deep layers having higher decoding compared to the other two with the superficial-deep difference decreasing with increasing mis-segmentation. In the middle depth signal case, only large misalignment values revert the sign of the middle and superficial layer decoding resulting in higher middle rather than superficial layer decoding accuracies. The initial middle layer accuracy was lower than the superficial layer accuracy (Fig. 1b, middle). When the signal originated from the superficial layers, the higher misalignment resulted in a decreased difference between superficial and middle layers. Taken together, layer mis-segmentation simulations show surprisingly stable decoding differences with larger changes only in high misalignment scenarios.

**Figure 5.**
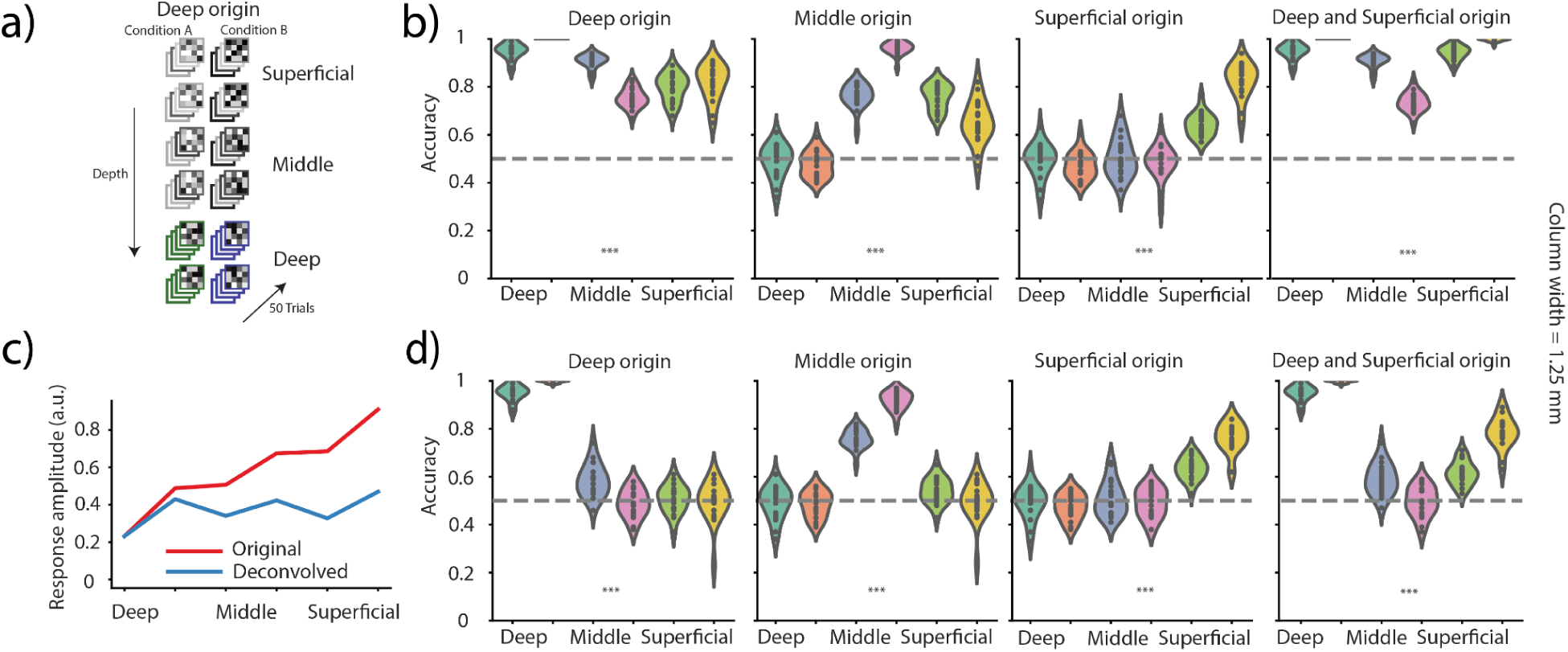
**Six layer simulations**. **a)** Deep origin simulation. Multivariate patterns were only consistent in the two deepest layers (green and blue), while other layers had different multivariate patterns in each of the 50 trials of each condition. **b)** Decoding accuracies across layers in deep, middle, superficial, and superficial and deep (same patterns) signal origin scenarios. Superficial and deep based on different patterns showed a similar pattern. **c)** Univariate response across layers of the original and deconvolved signals. **d)** Same as b) but the voxel responses were deconvolved using an inversion of the vascular model with the same parameters as the initial convolution. *** one-way ANOVA *p* < 0.001.

### Inaccurate deconvolution parameters do not greatly affect decoding results

As in the case of laminar delineation, the exact parameters needed for deconvolution of the laminar signal are usually unknown, as the exact parameter value would depend on the anatomical area (Markuerkiaga et al., 2016, 2021). We thus inverted the vascular model using parameters that differed from the parameters used in generating the data to simulate more realistic scenarios. Lower values of the parameter that controls the amount of draining expressed as percentages of the original remove a smaller extent of the laminar signal while larger values remove more. As in the mis-segmentation simulation, we plot t-values comparing the layer-of-interest to the other two layers (Fig. 4b).

In the case that the signal originates from either deep or middle layers, we see a decrease in the size of the effect between middle vs. deep and middle vs. superficial, respectively for smaller vascular parameters, which is driven by a smaller extent of signal removal from the more superficial layers in each case. In the middle origin scenario, we even see a reversal back to higher superficial rather than middle decoding accuracies, as seen in the non-deconvolved signal (Fig. 1c). The superficial signal origin simulation shows a trivial effect of a reducing superficial to middle and deep difference as a result of a larger vascular parameter.

### Simulation of six layers improves decoding accuracy

Finally, we examined whether increasing the number of simulated layers could result in a more precise decoding accuracy profile. This was motivated by prior methodological work suggesting that extracting more, thinner layers than the nominal voxel resolution allows can reduce partial volume effects and depth pooling, thereby sharpening laminar profiles (Huber, 2019; Huber et al., 2017, 2018). We used the same model, but increased the number of simulated layers to six. Deep, middle, and superficial depths thus correspond to two rather than one layer (Fig. 5a). We find increased precision in the decoding profile in both the non-deconvolved (Fig. 5b) and especially in the deconvolved (Fig. 5c) scenarios of deep, middle, superficial, and deep and superficial same pattern origin. This shows that increasing the number of layers might yield a more accurate reflection of the underlying signal.

**Figure 6:**
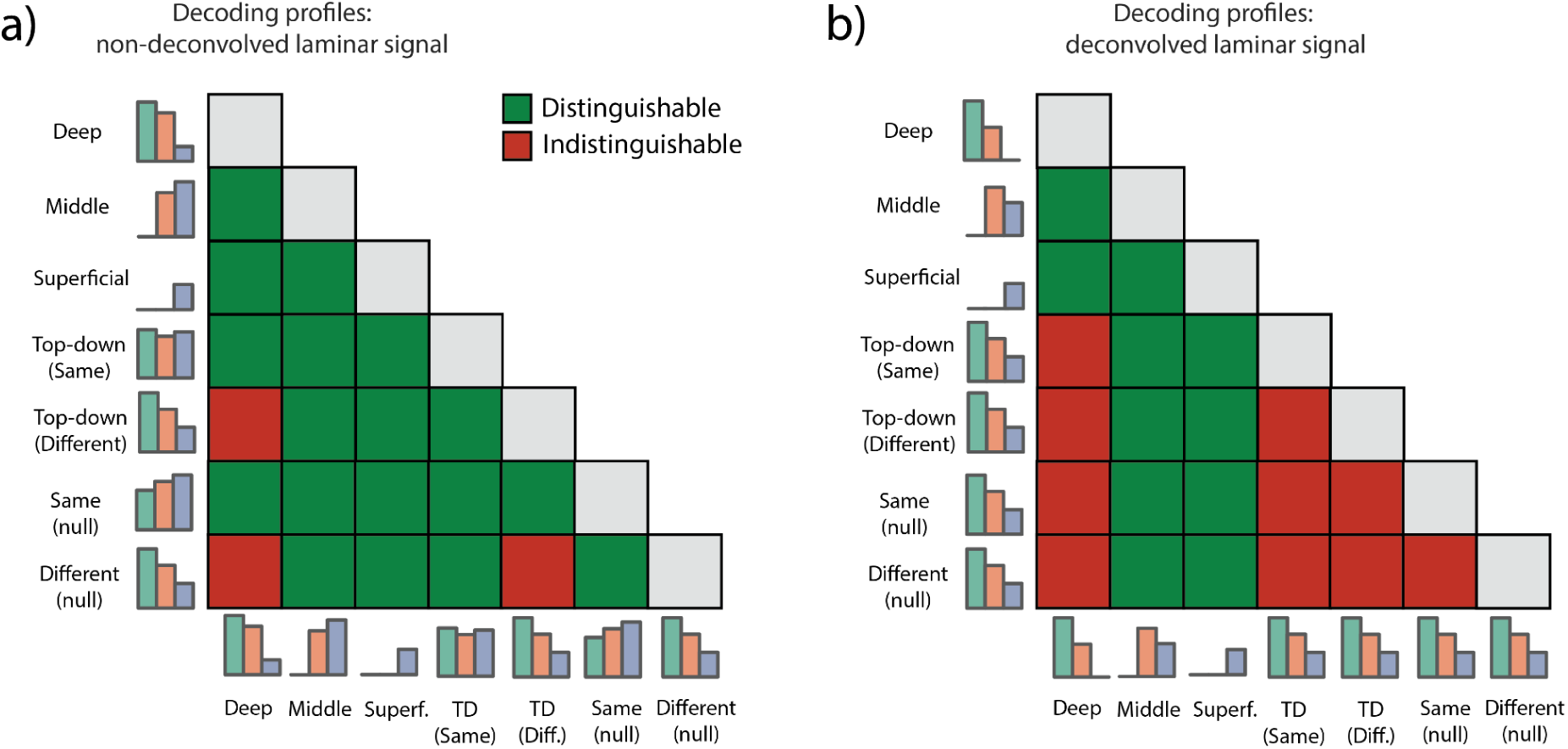
Qualitative comparison of laminar patterns based on signal origin in the three layer scenarios. **a)** A pairwise qualitative comparison of decoding accuracy profiles obtained based on non-deconvolved signals. Green indicates the ability to distinguish between two triplet laminar profiles, while red indicates an inability to do so. **b)** Same as a), but for decoding accuracies based on deconvolved signals.

## Discussion

In this study we investigated the impact of signal blurring due to draining veins on multivariate decoding accuracy across cortical layers. Multivariate decoding has become widely used to analyze high-resolution data (Bergmann et al., 2024; de Hollander et al., 2021; Degutis et al., 2024; Haenelt et al., 2023; Iamshchinina, Haenelt, et al., 2021; Iamshchinina, Kaiser, et al., 2021; Muckli et al., 2015; Ng et al., 2021; Thomas et al., 2024; Vizioli et al., 2020), as univariate analyses fail to capture the complexity of patterns in the data (Haxby et al., 2001; Haynes & Rees, 2006).

We simulated voxel responses across depth where the neuronal signal originated in either one of three layers, two layers, or in all layers. The simulated responses included a vascular draining model which blurred signals of more superficial layers by deeper layers. We find that in the single layer signal origin scenarios we can differentiate between layers, with the layer-of-origin having the highest decoding accuracy in all cases, but the middle origin scenario. The specificity of the decoding profiles was enhanced after deconvolving the signal. Inaccurate deconvolution parameters had little effect on these findings. In the two-layer scenarios which resembled a canonical top-down response, the accuracy of the decoding depended on whether the pairs of patterns present in superficial and deep layers were the same or different pairs; only the former scenario showed a profile matching the ground truth. Unlike the single origin simulations, the null scenarios yielded false positives; either the superficial or deep layer had the highest decoding accuracy, despite the expectation of no laminar differences. Finally, we saw an improvement in accuracy of the profiles when six layers were simulated rather than three.

Despite decoding accuracies of some scenarios not matching the ground truth, the triplet profiles could potentially serve as an identifier for an underlying signal. We thus qualitatively compare between the seven scenarios indicating whether the two triplet accuracy profiles can be distinguished from one another (Fig. 6). Most scenarios can be distinguished from one another when the signal was not deconvolved (Fig. 6a). We only encounter problems in distinguishing between the deep origin, different pattern top-down, and different pattern null scenario, in which the deep layer has the highest decoding accuracy.

Common amplitude-normalization or condition-contrast strategies (Aitken et al., 2020; de Hollander et al., 2021; Kok et al., 2016; Lawrence et al., 2018; Lawrence, Norris, et al., 2019) reduce superficial amplitude biases but do not undo pattern mixing. For example Lawrence et al. (2018, 2019) normalized each layer’s time course to control for amplitude differences, while another study has proposed using resting state fluctuation amplitudes for this purpose (Guidi et al., 2020). Yet, these approaches fail to account for the blurring of the signal in superficial layers. Voxel-wise regression from each adjacent deeper voxel, as previously applied to the entorhinal cortex (Koster et al., 2018), may remove neuronal responses shared between layers including physiological noise and therefore misattribute these signals to deeper layers. Multivariate analyses mitigate the blurring problem as classifiers penalize for high variance, so decoding accuracies will be lower for a mixture of laminar signals. We see this in our simulations with superficial layers having decreased decoding accuracies. Notably, our univariate activity of the superficial layer remains higher than middle and deep (Fig. 1c). This disentanglement between univariate and multivariate results has also been observed in other studies (Bergmann et al., 2024).

Based on our results, deconvolving the BOLD signal only works well when there is a single neural origin, yet it magnifies the false positive effect when neural signals originate from two or all depths or the cortex. A higher number of indistinguishable profiles pairs are present when the signal is deconvolved (Fig. 6b). In all problematic cases we see highest decoding in deep layers and decreasing decoding in middle and superficial layers. We are unable to differentiate between the two null, the two top-down, and the deep layer origin scenarios. With deconvolution the univariate superficial layer bias is replaced by a multivariate deep layer bias. One way to resolve these problems is to try to simulate responses by taking into account the point-spread function and vascular draining. However, such simulations might be challenging as parameters depend both on the region and individual subject neurophysiology.

In addition to examining the effects of draining, we investigated the influence of spatial size of the multivariate patterns on decoding accuracy. Since we kept the contrast-to-noise the same in all simulations when comparing the patterns, we see that the spatial size directly impacts decoding accuracy and interacts with the point-spread function of the layers. Previous studies have shown that the base decoding rate differs across regions of the cortex (Bhandari et al., 2018). For example the prefrontal cortex (PFC) is known to have lower decoding accuracies compared to visual areas. PFC neurons are separated laterally by 0.3 mm show tuning for visuospatial working memory information (Constantinidis et al., 2001), while large spatial biases might underlie the decoding of orientation-specific signals in visual areas (Beckett et al., 2012; Freeman et al., 2011). Thus, the BOLD signal in typical voxel sizes might not capture the information being encoded in the PFC as a consequence of several factors such as the microvasculature, signal-to-noise ratio, the heterogeneity of the stored information, and microanatomy (Bhandari et al., 2018; Goldman-Rakic, 1995; Rigotti et al., 2013). The latter directly relates to the spatial size of the patterns. The size of the spatial pattern across different layers might also differ as a result of an anatomical separation of population underlying different cognitive functions. For example the size of the multivariate patterns elicited by a motor response might differ compared to the patterns of stimulus maintenance (Degutis et al., 2024). This might create a problem directly comparing between decoding accuracies in different layers even within the same region across multiple cognitive task conditions, as for example examining an interaction effect between laminar activity and different cognitive conditions.

Simulating six layers rather than three layers improved the accuracy of the laminar accuracy profiles, especially on deconvolved signals. Draining together with the changes in noise and PSF results in an overall moderate shift in the location of highest decoding. Hence, the six layer scenario, unlike the three layer simulation, is less likely to mis-locate this moderate decoding shift and thus end up in the wrong depth bin. Multiple studies have used oversampled layers when conducting decoding analyses (Bergmann et al., 2024; Muckli et al., 2015). Extracting more layers than the number of voxels spanning the cortical depth can sharpen depth profiles, as the signal becomes less blurred and partial-voluming has a lesser effect. In our six layer simulations we see that the deep and superficial scenario on deconvolved signals shows no deep layer bias - as seen in the three layer scenario. Hence, deconvolution might work better when applied to oversampled laminar signals rather than only three depths.

One limitation in that the vascular draining model that we use does not generalize to all scenarios as it only characterizes signal blurring at steady state (Markuerkiaga et al., 2016, 2021); this might not be applicable to event-related measurements in which temporal dynamics of blood flow need to be considered (Havlicek & Uludağ, 2020; Uludag & Havlicek, 2021). Notably, we also do not explicitly simulate signals originating from pial veins on the cortical surface, even though these veins influence signals observed using GE-BOLD imaging (Goense et al., 2007; Goense & Logothetis, 2006; Koopmans et al., 2010, 2011; Polimeni et al., 2010). However, this pial vein blurring might be part of the differences reflected in the depth-dependent point-spread function. One could also substitute the draining vascular model for a more complex vascular model derived from first principles of the neuroanatomy and vascular circuitry of a given region (Báez-Yáñez et al., 2024; Bollmann et al., 2022; Uludag & Havlicek, 2021), yet an application of such a model to the scale of imaging analyses might yield similar results.

In summary, we show through simulations that multivariate decoding accuracies are influenced by draining veins in both single-layer signal origin scenarios and layer-unspecific signal scenarios. Additionally, our results insights into how the spatial scale of patterns influences multivariate accuracy. We hope this model provides the possibility to generate hypothesis-driven simulations that can be compared to experimental laminar results.

## Contributions

Conceptualization: J.K.D., D.C., R.L.; Methodology: J.K.D., D.C., R.L.; Formal Analysis: J.K.D.; Software: J.K.D., D.C.; Visualization: J.K.D.; Funding Acquisition: R.L.; Writing - Original Draft Preparation: J.K.D., R.L.; Writing – Review & Editing: J.K.D., D.C., R.L.; Supervision: R.L.

---

**Supplementary Table 1.** T- and *p*-values, respectively, corresponding to Figure 2b.

|  | Deep-middle | Deep-superficial | Middle-superficial |
| --- | --- | --- | --- |
| Non-deconvolved 1.25 mm. | 4.76<br>$p < 0.001$ | 5.27<br>$p < 0.001$ | 2.51<br>0.021 |
| Non-deconvolved 1 mm. | 6.95<br>$p < 0.001$ | 8.34<br>$p < 0.001$ | 0.87<br>0.39 |
| Non-deconvolved 0.83 mm. | 3.27<br>0.004 | 3.26<br>0.004 | 0.6<br>0.55 |
| Non-deconvolved 0.71 mm. | 0.39<br>0.70 | 0.067<br>0.95 | 0.31<br>0.75 |
| Deconvolved 1.25 mm. | 2.29<br>0.033 | 21.5<br>$p < 0.001$ | 11.3<br>$p < 0.001$ |
| Deconvolved 1 mm. | 3.72<br>0.0014 | 13.5<br>$p < 0.001$ | 6.59<br>$p < 0.001$ |
| Deconvolved 0.83 mm. | 2.98<br>0.0077 | 7.58<br>$p < 0.001$ | 2.91<br>0.0089 |
| Deconvolved 0.71 mm. | 1.21<br>0.24 | 2.86<br>0.01 | 0.88<br>0.39 |

**Supplementary Table 2.** T- and *p*-values, respectively, corresponding to Figure 2d.

|  | Deep-middle | Deep-superficial | Middle-superficial |
| --- | --- | --- | --- |
| Non-deconvolved 1.25 mm. | 5.54<br>$p < 0.001$ | 5.54<br>$p < 0.001$ | N/A |
| Non-deconvolved 1 mm. | 13.4<br>$p < 0.001$ | 14.2<br>$p < 0.001$ | 3.47<br>0.003 |
| Non-deconvolved 0.83 mm. | 9.52<br>$p < 0.001$ | 28.8<br>$p < 0.001$ | 8.52<br>$p < 0.001$ |
| Non-deconvolved 0.71 mm. | 4.09<br>$p < 0.001$ | 11.4<br>$p < 0.001$ | 6.89<br>$p < 0.001$ |
| Deconvolved 1.25 mm. | 1.02<br>0.32 | 2.09<br>0.05 | 0.69<br>0.49 |
| Deconvolved 1 mm. | 2.14<br>0.045 | 6.61<br>$p < 0.001$ | 2.95<br>0.0083 |
| Deconvolved 0.83 mm. | 3.95<br>$p < 0.001$ | 8.70<br>$p < 0.001$ | 3.0<br>0.007 |
| Deconvolved 0.71 mm. | 2.92<br>0.009 | 5.32<br>$p < 0.001$ | 1.87<br>0.076 |

**Supplementary Table 3.** T- and *p*-values, respectively, corresponding to Figure 2f.

|  | Deep-middle | Deep-superficial | Middle-superficial |
| --- | --- | --- | --- |
| Non-deconvolved 1.25 mm. | 1.94<br>0.068 | 5.01<br>$p<0.001$ | 3.58<br>0.002 |
| Non-deconvolved 1 mm. | 3.32<br>0.0035 | 8.61<br>$p<0.001$ | 4.13<br>$p<0.001$ |
| Non-deconvolved 0.83 mm. | 2.80<br>0.011 | 5.65<br>$p<0.001$ | 1.69<br>0.11 |
| Non-deconvolved 0.71 mm. | 1.79<br>0.089 | 1.56<br>0.13 | 0.15<br>0.88 |
| Deconvolved 1.25 mm. | 8.47<br>$p<0.001$ | 20.6<br>$p<0.001$ | 8.13<br>$p<0.001$ |
| Deconvolved 1 mm. | 13.1<br>$p<0.001$ | 20.3<br>$p<0.001$ | 4.81<br>$p<0.001$ |
| Deconvolved 0.83 mm. | 6.92<br>$p<0.001$ | 12.3<br>$p<0.001$ | 1.77<br>0.093 |
| Deconvolved 0.71 mm. | 2.01<br>0.058 | 3.62<br>0.0018 | 0.65<br>0.52 |

## References

Aitken, F., Menelaou, G., Warrington, O., Koolschijn, R. S., Corbin, N., Callaghan, M. F., & Kok, P. (2020). Prior expectations evoke stimulus-specific activity in the deep layers of the primary visual cortex. PLOS Biology, 18(12), e3001023. 10.1371/journal.pbio.3001023

Báez-Yáñez, M. G., Siero, J. C. W., Curcic, V., Van Osch, M. J. P., & Petridou, N. (2024). On the influence of the vascular architecture on Gradient Echo and Spin Echo BOLD fMRI signals across cortical depth: A simulation approach based on realistic 3D vascular networks. Neuroscience. 10.1101/2024.05.30.596593

Bastos, A. M., Loonis, R., Kornblith, S., Lundqvist, M., & Miller, E. K. (2018). Laminar recordings in frontal cortex suggest distinct layers for maintenance and control of working memory. Proceedings of the National Academy of Sciences, 115(5), 1117–1122. 10.1073/pnas.1710323115

Bastos, A. M., Usrey, W. M., Adams, R. A., Mangun, G. R., Fries, P., & Friston, K. J. (2012). Canonical Microcircuits for Predictive Coding. Neuron, 76(4), 695–711. 10.1016/j.neuron.2012.10.038

Beckett, A., Peirce, J. W., Sanchez-Panchuelo, R.-M., Francis, S., & Schluppeck, D. (2012). Contribution of large scale biases in decoding of direction-of-motion from high-resolution fMRI data in human early visual cortex. NeuroImage, 63(3), 1623–1632. 10.1016/j.neuroimage.2012.07.066

Bergmann, J., Petro, L. S., Abbatecola, C., Li, M. S., Morgan, A. T., & Muckli, L. (2024). Cortical depth profiles in primary visual cortex for illusory and imaginary experiences. Nature Communications, 15(1), 1002. 10.1038/s41467-024-45065-w

Bhandari, A., Gagne, C., & Badre, D. (2018). Just above Chance: Is It Harder to Decode Information from Prefrontal Cortex Hemodynamic Activity Patterns? Journal of Cognitive Neuroscience, 30(10), 1473–1498. 10.1162/jocn_a_01291

Bollmann, S., Mattern, H., Bernier, M., Robinson, S. D., Park, D., Speck, O., & Polimeni, J. R. (2022). Imaging of the pial arterial vasculature of the human brain in vivo using high-resolution 7T time-of-flight angiography. eLife, 11, e71186. 10.7554/eLife.71186

Chaimow, D., Degutis, J. K., Haenelt, D., Trampel, R., Weiskopf, N., & Lorenz, R. (2025). Challenges in replicating layer-specificity of working memory processes in human dlPFC. Neuroscience. 10.1101/2025.01.31.635930

Chaimow, D., Uğurbil, K., & Shmuel, A. (2018). Optimization of functional MRI for detection, decoding and high-resolution imaging of the response patterns of cortical columns. NeuroImage, 164, 67–99. 10.1016/j.neuroimage.2017.04.011

Chaimow, D., Yacoub, E., Uğurbil, K., & Shmuel, A. (2018). Spatial specificity of the functional MRI blood oxygenation response relative to neuronal activity. NeuroImage, 164, 32–47. 10.1016/j.neuroimage.2017.08.077

Constantinidis, C., Franowicz, M. N., & Goldman-Rakic, P. S. (2001). Coding Specificity in Cortical Microcircuits: A Multiple-Electrode Analysis of Primate Prefrontal Cortex. The Journal of Neuroscience, 21(10), 3646–3655. 10.1523/JNEUROSCI.21-10-03646.2001

de Hollander, G., van der Zwaag, W., Qian, C., Zhang, P., & Knapen, T. (2021). Ultra-high field fMRI reveals origins of feedforward and feedback activity within laminae of human ocular dominance columns. NeuroImage, 228, 117683. 10.1016/j.neuroimage.2020.117683

De Martino, F., Moerel, M., Ugurbil, K., Goebel, R., Yacoub, E., & Formisano, E. (2015). Frequency preference and attention effects across cortical depths in the human primary auditory cortex. Proceedings of the National Academy of Sciences, 112(52), 16036–16041. 10.1073/pnas.1507552112

Degutis, J. K., Chaimow, D., Haenelt, D., Assem, M., Duncan, J., Haynes, J.-D., Weiskopf, N., & Lorenz, R. (2024). Dynamic layer-specific processing in the prefrontal cortex during working memory. Communications Biology, 7(1). 10.1038/s42003-024-06780-8

Duvernoy, H. M., Delon, S., & Vannson, J. L. (1981). Cortical blood vessels of the human brain. Brain Research Bulletin, 7(5), 519–579. 10.1016/0361-9230(81)90007-1

Felleman, D. J., & Van Essen, D. C. (1991). Distributed Hierarchical Processing in the Primate Cerebral Cortex. Cerebral Cortex, 1(1), 1–47. 10.1093/cercor/1.1.1

Fracasso, A., Dumoulin, S. O., & Petridou, N. (2021). Point-spread function of the BOLD response across columns and cortical depth in human extra-striate cortex. Progress in Neurobiology, 202, 102034. 10.1016/j.pneurobio.2021.102034

Freeman, J., Brouwer, G. J., Heeger, D. J., & Merriam, E. P. (2011). Orientation Decoding Depends on Maps, Not Columns. The Journal of Neuroscience, 31(13), 4792–4804. 10.1523/JNEUROSCI.5160-10.2011

Goense, J. B. M., & Logothetis, N. K. (2006). Laminar specificity in monkey V1 using high-resolution SE-fMRI. Magnetic Resonance Imaging, 24(4), 381–392. 10.1016/j.mri.2005.12.032

Goense, J. B. M., Zappe, A.-C., & Logothetis, N. K. (2007). High-resolution fMRI of macaque V1. Magnetic Resonance Imaging, 25(6), 740–747. 10.1016/j.mri.2007.02.013

Goldman-Rakic, P. S. (1995). Cellular basis of working memory. Neuron, 14(3), 477–485. 10.1016/0896-6273(95)90304-6

Guidi, M., Huber, L., Lampe, L., Merola, A., Ihle, K., & Möller, H. E. (2020). Cortical laminar resting-state signal fluctuations scale with the hypercapnic blood oxygenation level-dependent response. Human Brain Mapping, 41(8), 2014–2027. 10.1002/hbm.24926

Haarsma, J., Kok, P., & Browning, M. (2022). The promise of layer-specific neuroimaging for testing predictive coding theories of psychosis. Schizophrenia Research, 245, 68–76. 10.1016/j.schres.2020.10.009

Haenelt, D., Chaimow, D., Nasr, S., Weiskopf, N., & Trampel, R. (2023). *Decoding of columnar-level organization across cortical depth using BOLD-and CBV-fMRI at 7 T* [Preprint]. Neuroscience. 10.1101/2023.09.28.560016

Havlicek, M., & Uludağ, K. (2020). A dynamical model of the laminar BOLD response. NeuroImage, 204, 116209. 10.1016/j.neuroimage.2019.116209

Haxby, J. V., Gobbini, M. I., Furey, M. L., Ishai, A., Schouten, J. L., & Pietrini, P. (2001). Distributed and Overlapping Representations of Faces and Objects in Ventral Temporal Cortex. Science, 293(5539), 2425–2430. 10.1126/science.1063736

Haynes, J.-D., & Rees, G. (2006). Decoding mental states from brain activity in humans. Nature Reviews Neuroscience, 7(7), 523–534. 10.1038/nrn1931

Heinzle, J., Koopmans, P. J., den Ouden, H. E. M., Raman, S., & Stephan, K. E. (2016). A hemodynamic model for layered BOLD signals. NeuroImage, 125, 556–570. 10.1016/j.neuroimage.2015.10.025

Huang, P., Correia, M. M., Rua, C., Rodgers, C. T., Henson, R. N., & Carlin, J. D. (2021). Correcting for Superficial Bias in 7T Gradient Echo fMRI. Frontiers in Neuroscience, 15, 715549. 10.3389/fnins.2021.715549

Huber, L. (2019, February 22). How many layers should I extract? Layer fMRI Blog. https://layerfmri.com/2019/02/22/how-many-layers-should-i-reconstruct/

Huber, L., Handwerker, D. A., Jangraw, D. C., Chen, G., Hall, A., Stüber, C., Gonzalez-Castillo, J., Ivanov, D., Marrett, S., Guidi, M., Goense, J., Poser, B. A., & Bandettini, P. A. (2017). High-Resolution CBV-fMRI Allows Mapping of Laminar Activity and Connectivity of Cortical Input and Output in Human M1. Neuron, 96(6), 1253–1263.e7. 10.1016/j.neuron.2017.11.005

Huber, L., Tse, D. H. Y., Wiggins, C. J., Uludağ, K., Kashyap, S., Jangraw, D. C., Bandettini, P. A., Poser, B. A., & Ivanov, D. (2018). Ultra-high resolution blood volume fMRI and BOLD fMRI in humans at 9.4 T: Capabilities and challenges. NeuroImage, 178, 769–779. 10.1016/j.neuroimage.2018.06.025

Huber, L., Uludağ, K., & Möller, H. E. (2019). Non-BOLD contrast for laminar fMRI in humans: CBF, CBV, and CMRO2. NeuroImage, 197, 742–760. 10.1016/j.neuroimage.2017.07.041

Iamshchinina, P., Haenelt, D., Trampel, R., Weiskopf, N., Kaiser, D., & Cichy, R. M. (2021). *Benchmarking GE-BOLD, SE-BOLD, and SS-SI-VASO sequences for depth-dependent separation of feedforward and feedback signals in high-field MRI* [Preprint]. Neuroscience. 10.1101/2021.12.10.472064

Iamshchinina, P., Kaiser, D., Yakupov, R., Haenelt, D., Sciarra, A., Mattern, H., Luesebrink, F., Duezel, E., Speck, O., Weiskopf, N., & Cichy, R. M. (2021). Perceived and mentally rotated contents are differentially represented in cortical depth of V1. Communications Biology, 4(1), 1069. 10.1038/s42003-021-02582-4

Kok, P., Bains, L. J., van Mourik, T., Norris, D. G., & de Lange, F. P. (2016). Selective Activation of the Deep Layers of the Human Primary Visual Cortex by Top-Down Feedback. Current Biology, 26(3), 371–376. 10.1016/j.cub.2015.12.038

Koopmans, P. J., Barth, M., & Norris, D. G. (2010). Layer-specific BOLD activation in human V1. Human Brain Mapping, 31(9), 1297–1304. 10.1002/hbm.20936

Koopmans, P. J., Barth, M., Orzada, S., & Norris, D. G. (2011). Multi-echo fMRI of the cortical laminae in humans at 7T. NeuroImage, 56(3), 1276–1285. 10.1016/j.neuroimage.2011.02.042

Koster, R., Chadwick, M. J., Chen, Y., Berron, D., Banino, A., Düzel, E., Hassabis, D., & Kumaran, D. (2018). Big-Loop Recurrence within the Hippocampal System Supports Integration of Information across Episodes. Neuron, 99(6), 1342–1354.e6. 10.1016/j.neuron.2018.08.009

Lawrence, S. J. D., Formisano, E., Muckli, L., & de Lange, F. P. (2019). Laminar fMRI: Applications for cognitive neuroscience. NeuroImage, 197, 785–791. 10.1016/j.neuroimage.2017.07.004

Lawrence, S. J. D., Norris, D. G., & de Lange, F. P. (2019). Dissociable laminar profiles of concurrent bottom-up and top-down modulation in the human visual cortex. eLife, 8, e44422. 10.7554/eLife.44422

Lawrence, S. J. D., van Mourik, T., Kok, P., Koopmans, P. J., Norris, D. G., & de Lange, F. P. (2018). Laminar Organization of Working Memory Signals in Human Visual Cortex. Current Biology, 28(21), 3435–3440.e4. 10.1016/j.cub.2018.08.043

Markuerkiaga, I., Barth, M., & Norris, D. G. (2016). A cortical vascular model for examining the specificity of the laminar BOLD signal. NeuroImage, 132, 491–498. 10.1016/j.neuroimage.2016.02.073

Markuerkiaga, I., Marques, J. P., Gallagher, T. E., & Norris, D. G. (2021). Estimation of laminar BOLD activation profiles using deconvolution with a physiological point spread function. Journal of Neuroscience Methods, 353, 109095. 10.1016/j.jneumeth.2021.109095

Marquardt, I., Schneider, M., Gulban, O. F., Ivanov, D., & Uludağ, K. (2018). Cortical depth profiles of luminance contrast responses in human V1 and V2 using 7 T fMRI. Human Brain Mapping, 39(7), 2812–2827. 10.1002/hbm.24042

Mendoza-Halliday, D., Major, A. J., Lee, N., Lichtenfeld, M. J., Carlson, B., Mitchell, B., Meng, P. D., Xiong, Y. (Sophy), Westerberg, J. A., Jia, X., Johnston, K. D., Selvanayagam, J., Everling, S., Maier, A., Desimone, R., Miller, E. K., & Bastos, A. M. (2024). A ubiquitous spectrolaminar motif of local field potential power across the primate cortex. Nature Neuroscience, 27(3), 547–560. 10.1038/s41593-023-01554-7

Muckli, L., De Martino, F., Vizioli, L., Petro, L. S., Smith, F. W., Ugurbil, K., Goebel, R., & Yacoub, E. (2015). Contextual Feedback to Superficial Layers of V1. Current Biology, 25(20), 2690–2695. 10.1016/j.cub.2015.08.057

Ng, A. K. T., Jia, K., Goncalves, N. R., Zamboni, E., Kemper, V. G., Goebel, R., Welchman, A. E., & Kourtzi, Z. (2021). Ultra-High-Field Neuroimaging Reveals Fine-Scale Processing for 3D Perception. The Journal of Neuroscience, 41(40), 8362–8374. 10.1523/JNEUROSCI.0065-21.2021

Norris, D. G. (2012). Spin-echo fMRI: The poor relation? NeuroImage, 62(2), 1109–1115. 10.1016/j.neuroimage.2012.01.003

Polimeni, J. R., Fischl, B., Greve, D. N., & Wald, L. L. (2010). Laminar analysis of 7T BOLD using an imposed spatial activation pattern in human V1. NeuroImage, 52(4), 1334–1346. 10.1016/j.neuroimage.2010.05.005

Ramadan, D., Mueller, S., Stirnberg, R., Bosch, D., Ehses, P., Scheffler, K., & Bause, J. (2025). Macrovascular contributions to resting-state fMRI signals: A comparison between EPI and bSSFP at 9.4 Tesla. Imaging Neuroscience, 3, imag_a_00435. 10.1162/imag_a_00435

Rigotti, M., Barak, O., Warden, M. R., Wang, X.-J., Daw, N. D., Miller, E. K., & Fusi, S. (2013). The importance of mixed selectivity in complex cognitive tasks. Nature, 497(7451), Article 7451. 10.1038/nature12160

Rojer, A. S., & Schwartz, E. L. (1990). Cat and monkey cortical columnar patterns modeled by bandpass-filtered 2D white noise. Biological Cybernetics, 62(5), 381–391. 10.1007/BF00197644

Sharoh, D., van Mourik, T., Bains, L. J., Segaert, K., Weber, K., Hagoort, P., & Norris, D. G. (2019). Laminar specific fMRI reveals directed interactions in distributed networks during language processing. Proceedings of the National Academy of Sciences, 116(42), 21185–21190. 10.1073/pnas.1907858116

Shmuel, A., Yacoub, E., Chaimow, D., Logothetis, N. K., & Ugurbil, K. (2007). Spatio-temporal point-spread function of fMRI signal in human gray matter at 7 Tesla. NeuroImage, 35(2), 539–552. 10.1016/j.neuroimage.2006.12.030

Thomas, E. R., Haarsma, J., Nicholson, J., Yon, D., Kok, P., & Press, C. (2024). Predictions and errors are distinctly represented across V1 layers. Current Biology, 34(10), 2265–2271.e4. 10.1016/j.cub.2024.04.036

Triantafyllou, C., Hoge, R. D., Krueger, G., Wiggins, C. J., Potthast, A., Wiggins, G. C., & Wald, L. L. (2005). Comparison of physiological noise at 1.5 T, 3 T and 7 T and optimization of fMRI acquisition parameters. NeuroImage, 26(1), 243–250. 10.1016/j.neuroimage.2005.01.007

Uludag, K., & Havlicek, M. (2021). Determining laminar neuronal activity from BOLD fMRI using a generative model. Progress in Neurobiology, 207, 102055. 10.1016/j.pneurobio.2021.102055

van Kerkoerle, T., Self, M. W., & Roelfsema, P. R. (2017). Layer-specificity in the effects of attention and working memory on activity in primary visual cortex. Nature Communications, 8(1), 13804. 10.1038/ncomms13804

Vizioli, L., De Martino, F., Petro, L. S., Kersten, D., Ugurbil, K., Yacoub, E., & Muckli, L. (2020). Multivoxel Pattern of Blood Oxygen Level Dependent Activity can be sensitive to stimulus specific fine scale responses. Scientific Reports, 10(1), 7565. 10.1038/s41598-020-64044-x

Yang, J., Huber, L., Yu, Y., & Bandettini, P. A. (2021). Linking cortical circuit models to human cognition with laminar fMRI. Neuroscience & Biobehavioral Reviews, 128, 467–478. 10.1016/j.neubiorev.2021.07.005

